# Predicting the phenotype of Mendelian disease missense mutations using amino acid conservation and protein stability change

**DOI:** 10.1101/086470

**Authors:** Maria T. Chavez, Ethan O. Perlstein

## Abstract

Many Mendelian diseases are caused by recessive, loss-of-function missense mutations. On a gene-by-gene basis, it has been demonstrated that missense mutations cause, among other defects, protein misfolding, protein instability, protein mistransport, which strongly suggests that pathogenic missense mutations do not occur at random positions. Based on those observations, we predicted that Mendelian disease missense mutations are enriched in evolutionarily-conserved amino acids. In a pilot set of 260 Mendelian diseases genes affecting cellular organelles we show that missense mutations indeed occur in amino acids that are significantly more conserved than the average amino acid in the protein based on three different scoring methods (Jensen Shannon Divergence p = 7.78E-03, Shannon Entropy p = 1.68E-13, Sum of Pairs p = 1.55E-17). In order to understand how these results might be related to clinical phenotypes in humans or preclinical phenotypes in model organisms, we calculated the protein stability change upon mutation (ΔΔGu) using EASE-MM and found that, on average, pathogenic mutations cause a stability change of greater magnitude than benign mutations (p = 4.414428E-23). Finally, we performed a computational case study on NPC1, the gene responsible for 95% of diagnosed cases of the lysosomal storage disorder Niemann-Pick Type C using a set of 411 missense mutations from the Exome Aggregation Consortium.

## Introduction

While individually rare diseases represent a disproportionately small fraction of the global patient population, there are over 7,000 Mendelian (single-gene) phenotypes reported in OMIM, representing in aggregate about 0.4% of live births (and about 8% of live births have a genetic disorder recognizable by early adulthood if all congenital abnormalities are included).^1^^,^^2^ Research into Mendelian diseases is essential not only for the treatment of their specific patient populations but also to widen our understanding of fundamental genes and pathways, which also affect common diseases^3^. For example, Mendelian forms of high and low blood pressure are caused by mutations that affect renal salt re-absorption and net salt balance; these discoveries identified promising therapeutic targets, such as KCNJ1, and provided the scientific basis for public health efforts to reduce heart attacks and strokes by reduction in dietary salt intake.^4^ Other common disease drugs based on Mendelian disease genetics include orexin antagonists for sleep^5^, BACE1 inhibitors for Alzheimer’s^6^, and PCSK9 inhibitors for cardiovascular disease and cholesterol management^7^.

Mendelian diseases provide a unique opportunity. Given that they are associated with a single gene, if this gene is evolutionarily conserved and present in simpler model organisms (e.g.s, *Caenorhadbitis elegans, Drosophila melanogaster, Danio rerio* and Saccharomyces cerevisiae), an organism-based disease model can be used to gain fundamental insights into human disease pathophysiology. Niemann-Pick Type C Disease (NPC) and Cystic Fibrosis (CF) are two examples of diseases where an evolutionary approach has been employed and proved to be fruitful. Yeast^8^, nematode^9^, fly^10^ and zebrafish^11^models of NPC have been developed and shed light into the role cholesterol storage and mobilization play in the disease. Similarly yeast^12^, nematodes^13^, flies^14^, and zebrafish^15^ models of CF have been developed and provided therapeutic target insights most notably, using a yeast model of CF Veit *et al* showed that the protein misfolding defect caused by the most common CF mutation, ΔF508, can be suppressed by an evolutionarily conserved suppressor gene RPL12.

While model organisms can significantly reduce the cost and time involved in the identification of key disease processes and novel therapies, a challenge remains. Which mutations should be used to generate novel disease models when dozens if not hundreds of mutations can cause the same Mendelian disease? How can we increase the likelihood of observing the same phenotypes in disease models that are observed in human patients remains an open question. We therefore sought to determine whether we could combine publicly available protein sequence data with publicly available protein stability prediction tools to predict the clinical and biochemical consequences of a pilot set of 411 missense mutations in the NPC1 gene from the Exome Aggregation Consortium database (ExAC), which contains highly-quality whole exomes from more than 60,000 unrelated individuals from around the world.

Previous attempts to predict the functional consequences -- if any -- of a given protein-coding mutation have been published. Conservation analyses are one of the most common approaches, and they have been used to detect residues involved in ligand binding^16^^,^^17^, protein-protein interaction interfaces (PPIs)^18^^,^^19^, maintaining structure^20^^,^^21^^,^^22^ and protein functional specificity^23^^,^^24^^,^^25^. Computational methods that don’t use conservation are generally only used in cases where there are no homologs and they usually work by either identifying local shared structural patterns or residues with unusual electrostatic and ioinzation properties^26^. Petrova and Wu showed that conservation is the most powerful attribute for predicting functional importance in the aforementioned examples^27^. However, conservation alone is not sufficient to predict all residues in PPIs^28^. While conservation is a common approach, there isn’t an agreed-upon method for scoring conservation. We therefore used three different methods with different mathematical properties (Jensen-Shannon Divergence, Shannon Entropy and Sum of Pairs) in order to score conservation. Additionally, it has been shown that while the value of ΔΔGu alone is not enough to reliably predict the clinical effect of a given single nucleotide variant, it can be combined with other methods to improve the characterization of pathogenic mutations^29^^,^^30^^,^^31^.

The two most commonly used methods to predict the functional effect of a given mutation are PolyPhen2 and SIFT. PolyPhen2 uses eight sequence-based and three structure based methods (profile-based scores, identity-based scores, MSA depth, residue volume, accessible surface area, hydrophobic propensity, B-factor) and the functional importance is then predicted from the individual features by a Bayes classifier^32^. SIFT, on the other hand, compiles a dataset of functionally related protein sequences using PSI-BLAST; builds an alignment; scans each position; calculates the probability of each of the 20 possible amino acids at that position; normalizes it by the probability of the most frequent amino acid; and finally predicts a mutation to affect protein function if the probability is below a threshold^33^^,^^34^^,^^35^.

Our analysis shows that Mendelian disease missense mutations occur in significantly conserved amino acids, and that pathogenic mutations are characterized by a stability change of greater magnitude, consistent with the findings of Folkman *et al^35^* and Casadio *et al^36^*. We found a positive correlation between the predicted ΔΔGu and the conservation score, which was stronger than the correlation between the computed relative accessible surface area and the conservation score. Additionally, comparing our predictions with those of PolyPhen2 and SIFT it appears, at least in the case of the lysosomal storage disorder gene NPC1, that our predictions are slightly more accurate than PolyPhen2’s and comparable to those of SIFT.

## Results

For this analysis we began with a list of 260 Mendelian disease genes found in Antonella De Matteis’ website (which can be found here: http://edit.tigem.it/en/research/researchers/antonella-de-matteis/disease-genes) These genes are involved in organellar biology, e.g., lysosomal storage disorders. For every gene for which an ortholog exists, we used Homologene to download the protein sequences from human (*Homo sapiens*), mouse (*Mus musculus*), fly (*Drosophila melanogaster*), zebrafish (*Danio rerio*), nematode (*Caenorhabditis elegans*) and yeast (*Saccharomyces cerevisiae*). If a sequence was missing we checked on OMA browser as well.

We then created a multiple sequence alignment using CLUSTAL W2. The second half of the dataset was aligned using CLUSTAL OMEGA, given that CLUSTAL W2 was deprecated during the summer of 2015. We then used Princeton’s Protein Residue Conservation Prediction tool to calculate three scoring methods for all protein sequences in the dataset. The first is Jensen Shannon Divergence, which quantifies the similarity between probability distributions using a background amino acid distribution. Positions in an alignment with distributions that are very different from the background distribution are then likely to be functionally important. The second is Shannon Entropy, one of the most common measures of conservation at a site, and which doesn’t take into account physicochemical similarity between amino acids. The Shannon Entropy score is smaller for fully conserved columns. The third is Sum of Pairs, which scores conservation using a similarity matrix and computes the pairwise similarity between all amino acids in a column. For consistency, the Shannon Entropy scores were scaled to the range [0,1] and subtracted from one. In methods where a background amino acid distribution or similarity matrices were required, we used BLOSUM 62. We then searched ClinVar1 (a public archive showing the relationship between human variants and observed phenotypes) for mutations associated with the gene of interest, and plugged this information into a program we wrote (which can be found on GitHub https://github.com/materechm/plabData/blob/master/scripts/conservation_analysis.py).

In the 260-gene dataset, the average number of mutations per gene is 11.6, with 19% of those being benign. On average, 5.24 fully conserved residues per protein are affected by pathogenic mutations; these residues comprise 3.7% of all fully conserved amino acids in the dataset. As shown in **Figure 1**, pathogenic mutations are, on average, more conserved than the average amino acid in the protein (Jensen Shannon Divergence p = 7.78E-03, Shannon Entropy p = 1.68E-13, Sum of Pairs p = 1.55E-17), and more conserved than benign mutations (Jensen Shannon Divergence p = 1.73E-05, Shannon Entropy p = 1.05E-26, Sum of Pairs p = 2.98E-27).

**Figure 1:**
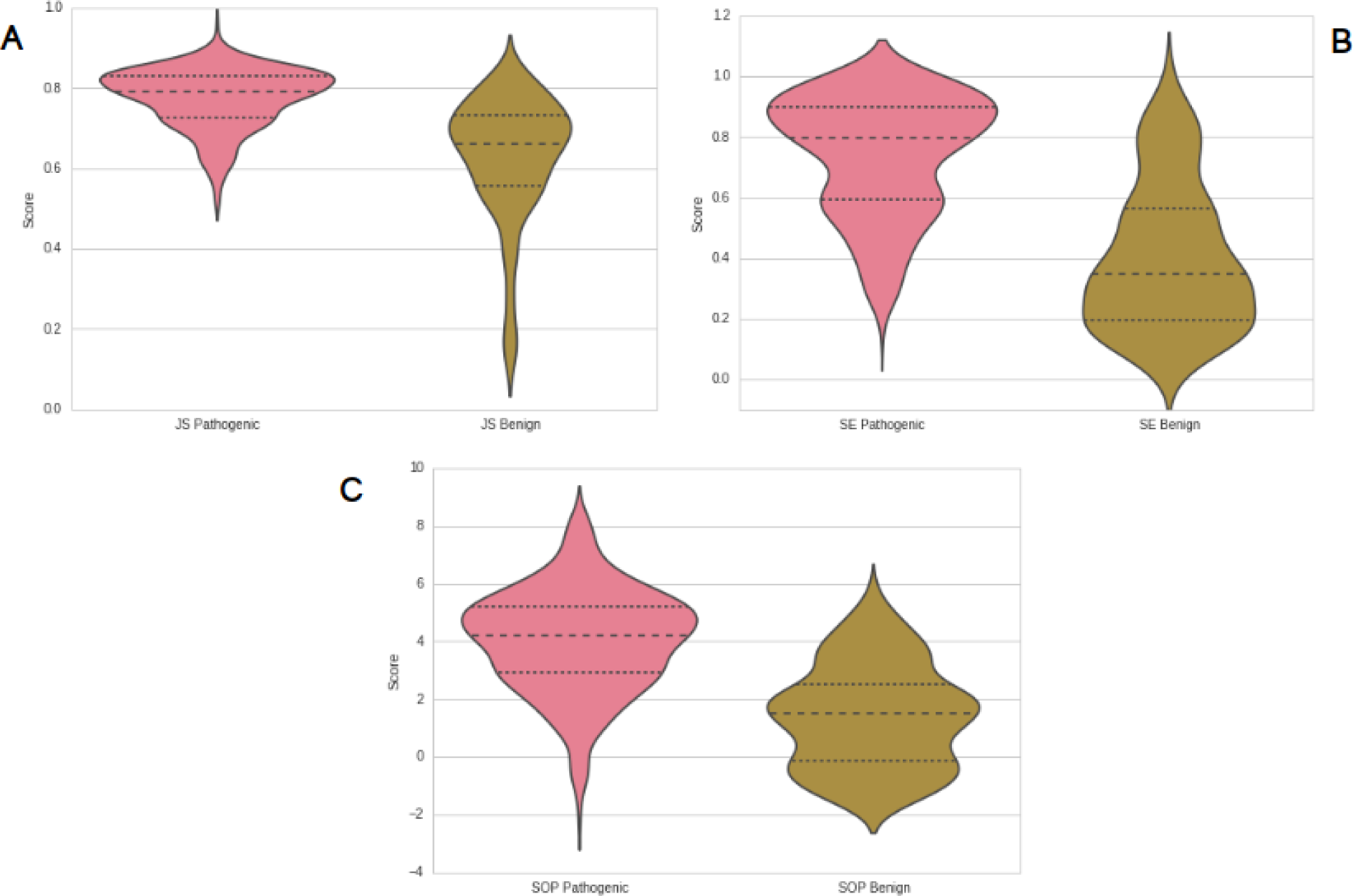
violin plots showing conservation scores for the set of pathogenic mutations (containing mutations found in the pilot 260 Mendelian disease genes) and benign mutations (containing mutations pilot 260 Mendelian disease genes). (A) shows Jensen-Shannon divergence scores, (B) shows Shannon Entropy scores, (C) shows Sum of Pairs scores. In all three sub-figures the pink violin plot represents the pathogenic scores and the beige plot represents the benign scores.

We performed a number of controls to ensure correct interpretation of the dataset. First, the difference in conservation between pathogenic mutations and the average amino acid seems to be more significant when the total number of mutations is higher, and the relationship (Pearson correlation = −0.361300) appears to follow a power trend (**Figure 2**), which means we get a stronger signal as mutation number increases, as expected.

**Figure 2:**
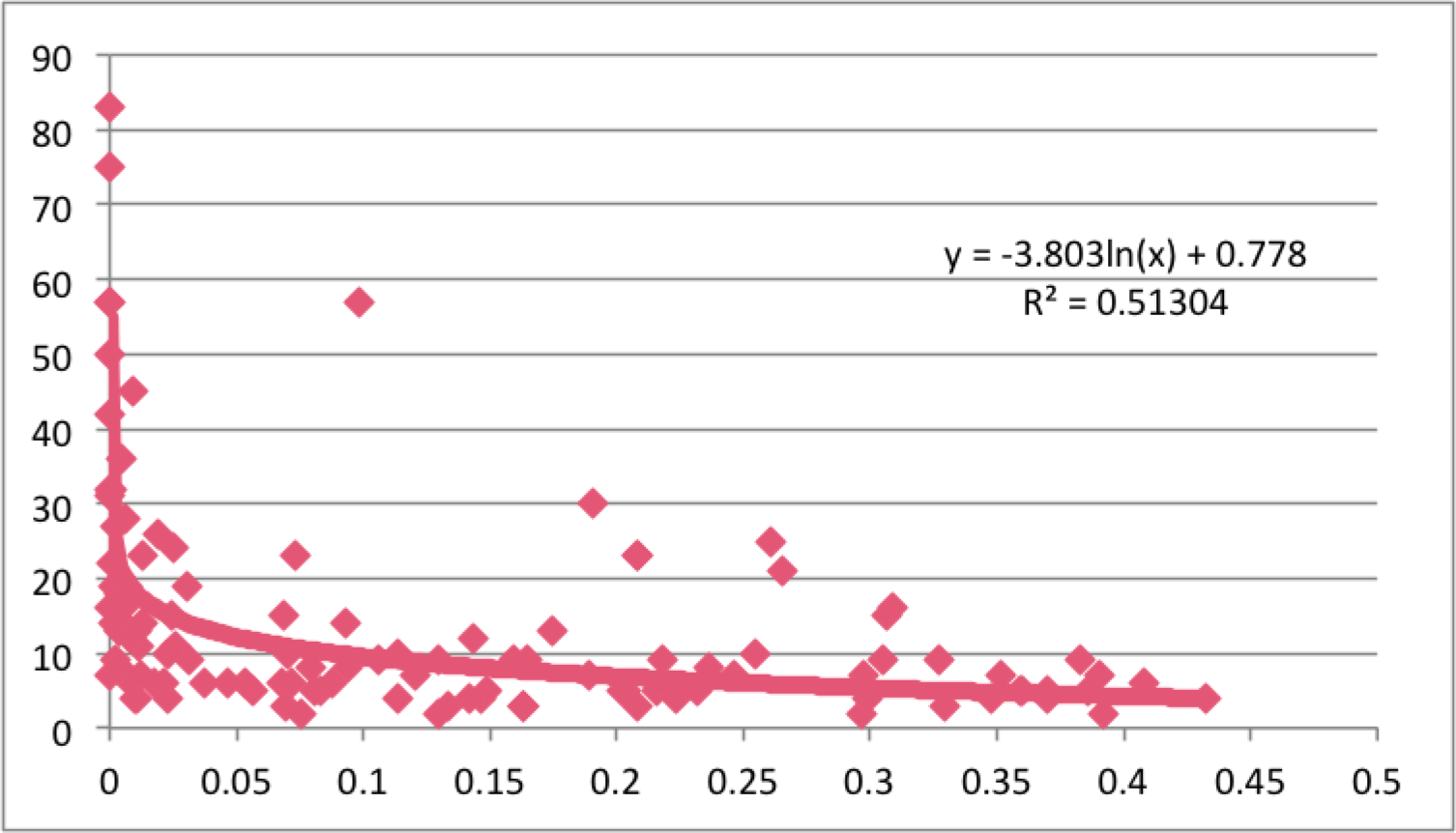
Graph showing the correlation between number of mutations (y axis) and p value (where < 0.05 is significant)

Second, there does not seem to be a relationship between the number of mutations and the percentage of fully conserved mutations (**Supplementary Materials/Figures**). This implies that even though the difference in conservation between pathogenic mutations and the average amino acid is more significant as mutation number increases, the percentage of fully conserved mutations does not depend on mutation number.

Third, there does not seem to be a correlation between the percentage fully conserved mutations or the p value with the number of species that have a disease gene homolog (**Supplementary Materials/Figures**), which implies that we aren’t observing a stronger signal just because a disease gene is incompletely conserved. In other words, the signal we observe doesn’t come from disease gene homologs that are only present in two or three (versus four or five) species.

To see if any further trends would appear in subsets of the data, we used the PANTHER (protein annotation through evolutionary relationship) classification system^37^^,^^38^ which groups genes according to their function using Gene Ontology (GO)^39^^,^^40^ annotations and performs an enrichment analysis showing which biological processes, cellular components and molecular functions are over (or under) represented. We found 35 molecular functions that are significantly overrepresented in the dataset (p < 0.05), and 6 molecular functions that are significantly underrepresented in the dataset (p < 0.05). Every function follows the trend we see in the entire dataset, where amino acid residues altered by pathogenic mutations are more conserved than the average amino acid in a protein. In 28 of these molecular functions, the difference in conservation between amino acid residues altered by pathogenic mutations and the average amino acid in the protein is statistically significant for at least one of the three scoring methods (p < 0.05). In 26 of those 28, the difference between the average scores in the subsets and the average scores in the dataset is statistically significant for at least one of the three scoring methods (p < 0.05).

The subsets of genes associated with NADH dehydrogenase (quinone) activity, UDP-glucosyltransferase activity, hydrolase activity, hydrolyzing O-glycosyl compounds, procollagen-lysine 5-dioxygenase activity, hydrolase activity acting on acid acting on acid anhydrides in phosphorus-containing anhydrides, hydrolase activity acting on acid anhydrides all have averages lower than the entire dataset (p < 0.05 for at least one scoring method) even though they do follow the trend we see with the entire dataset.

The subset of genes associated with receptor activity, DNA binding and molecular transducer activity have an average higher than the entire dataset (p < 0.05 for at least one scoring method) and the subset of genes associated with signaling receptor activity and nucleic acid binding have an average lower than the entire dataset (p < 0.05 for at least one scoring method). All of these are significantly underrepresented in the dataset and have a small number of genes (2-7) so it is likely the samples are too small to make any valid conclusions.

To see if we could get any further information about both clinical significance and the likelihood of a mutation producing a phenotype in an animal model, we used EASE-MM5 (Evolutionary, Amino acid, and Structural Encodings with Multiple Models), which uses five specialized support vector machines (SVMs) to predict protein stability changes (ΔΔGu) based on predicted secondary structures (SS) and relative accessible surface area (RSA) of mutated residues. This data can be found in the protein data spreadsheet on Github. The values on the stability change column can be interpreted as follows: values from -infinity to −1 indicate the mutation is destabilizing, from −1 to −0.5 indicate that the mutation is likely destabilizing, from −0.5 to 0.5 that they are likely stabilizing and from 0.5 to infinity indicate that they are stabilizing. The possible secondary structures are helix (H), extended/sheet (E), or coil (C). Finally, the relative accessible surface area is expressed as a percentage (in decimal form).

### Case Analysis: Gene Clusters

We initially performed the protein stability analysis on eight of the functional subsets generated by PANTHER. Mutations resulting in deletions and or a premature stop codon were excluded from the analysis. Below is the data for the top two subsets based on number of genes and p values plus the data for the combination of all eight subsets. The rest of the data can be found in Supplementary data/materials.

In the subset associated with hydrolase activity hydrolyzing O-glycosyl compounds we have 16 genes, including HEXA (which is associated with Tay-Sachs disease), containing 218 pathogenic mutations and 46 benign mutations total after eliminating amino acid deletions and mutations resulting in stop codons. We see the same trends in the conservation analysis for this subset than we see on the conservation analysis performed in the entire dataset: throughout all methods pathogenic mutations are more conserved than benign mutations (Jensen Shannon Divergence average: 0.778107235, 0.6591219565 respectively p = 3.37E-07, Shannon Entropy average: 0.7303362673, 0.4231956522 respectively p = 5.32E-10, Sum of Pairs average: 4.03976, 1.5829304348 p = 6.38E-11). Note: given that the current protein analysis only includes pathogenic and benign mutation scores, figures and data comparing pathogenic mutations versus the average amino acid in the protein are not included (**Figure 3A-C**).

**Figure 3:**
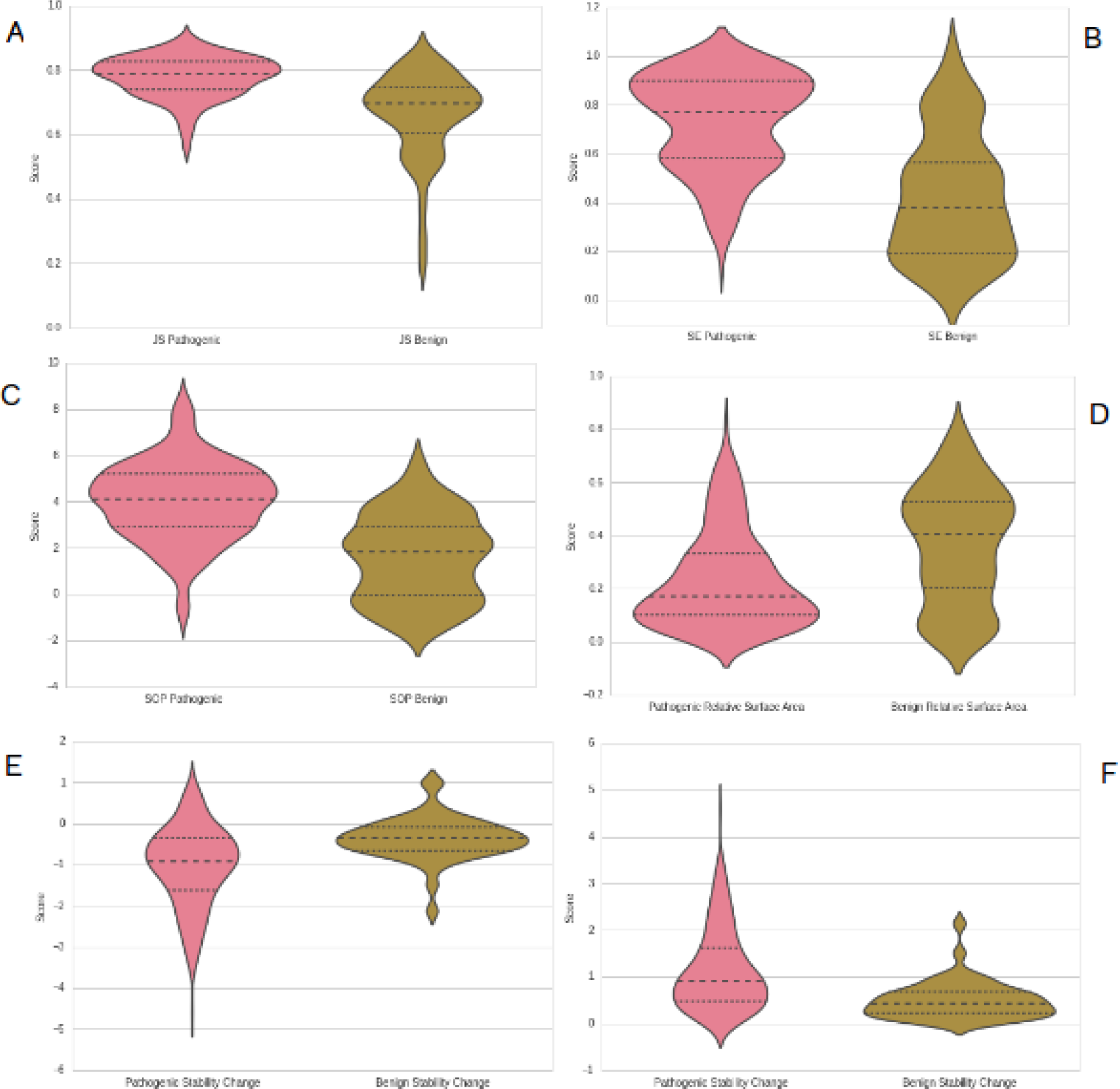
violin plots for the PANTHER subset hydrolase activity hydrolyzing O-glycosyl compounds showing (A) Jensen-Shannon Divergence score (B) Shannon Entropy score (C) Sum of Pairs score (D) Relative accessible surface area (E) ΔΔGu (F) absolute value (magnitude) of ΔΔGu. In all six sub-figures the pink violin plot represents the pathogenic scores and the beige plot represents the benign scores.

Pathogenic mutations are on average more destabilising than benign mutations (average stability: −1.039743578, 0.9785293086 respectively, p = 2.335294E-08). We also see a greater decrease in relative accessible surface area caused by pathogenic mutations than benign mutations (average relative accessible surface area: 0.2277522936, 0.362826087 respectively, p = 0.0001645118, **Figure 3D,E**).

We also see a negative correlation between the stability score and conservation score with all three scoring methods: Jensen Shannon divergence, Shannon Entropy and Sum of Pairs (Pearson correlation coefficient −0.356210, −0.302972 and −0.395452 respectively) and a negative correlation between relative accessible surface area and conservation score with all 3 scoring methods (Pearson correlation coefficient: −0.473328, −0.500113 and −0.448384 respectively) in this subset and there is a positive correlation between the stability score and relative accessible surface area (Pearson correlation coefficient: 0.480174).

Casadio *et al* showed that the correlation of disease probability and protein stability change is greater using the magnitude of experimentally measured stability change rather than the stability change itself, given that mutations with a high stability score might have had stabilization via loss of flexibility (which can lead to loss of function). Therefore we decided to redo the stability analysis using the absolute value of the computed ΔΔGu.

Taking the absolute value the average score for pathogenic mutations is 1.1381059633 and the average score for benign mutations is 0.4991782609 (p = 5.68115E-10). We also see a positive correlation between the stability score and conservation score with all three scoring methods: Jensen Shannon divergence, Shannon Entropy and Sum of Pairs (Pearson correlation coefficient: 0.340070, 0.297829 and 0.398375 respectively) which suggests that the magnitude of the stability change is greater the more conserved the amino acid is (**Figure 3F**). We see a negative correlation between the stability change and the relative accessible surface area, though the correlation is weaker than in the previous analysis (Pearson correlation coefficient: −0.446044).

In the subset associated with mannosyltransferase activity we have 11 genes, including POMT1 (which is associated with Walker-Warburg Syndrome), containing 51 pathogenic mutations and 20 benign mutations total after eliminating amino acid deletions and mutations resulting in stop codons. We see the same trends in the conservation analysis for this subset than we see on the conservation analysis performed in the entire dataset: throughout all methods pathogenic mutations are more conserved than benign mutations (Jensen Shannon Divergence average: 0.7410958824, 0.5645155 respectively p = 0.0007190673, Shannon Entropy average: 0.6749821569, 0.391811 respectively p = 8.25E-05, Sum of Pairs average: 3.3076778431, 1.1831355 p = 0.0001877878). Note: given that the current protein analysis only includes pathogenic and benign mutation scores, figures and data comparing pathogenic mutations versus the average amino acid in the protein are not included (**Figure 4A-C**))

**Figure 4:**
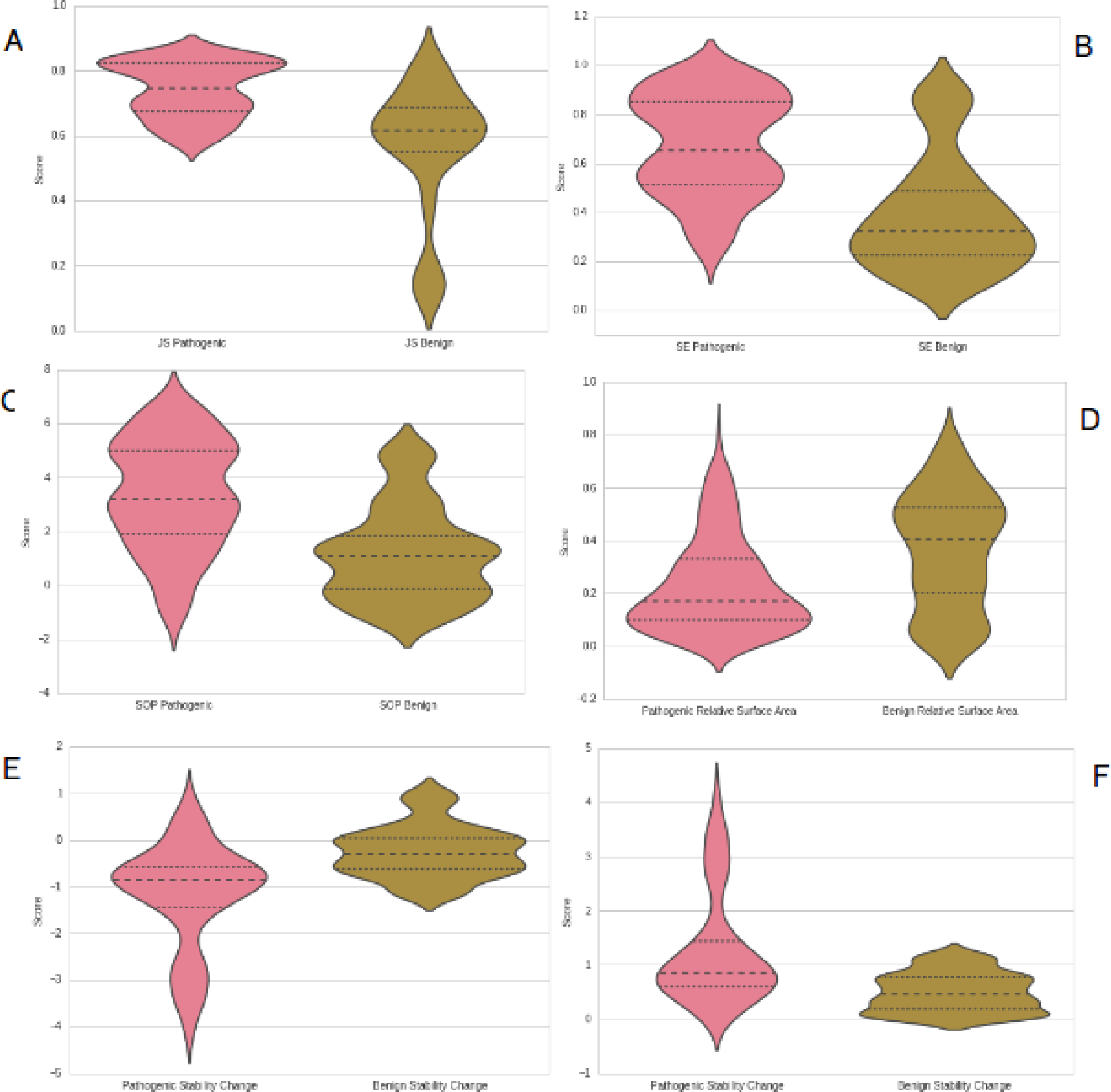
violin plots for the PANTHER subset mannosyltransferase activity showing (A) Jensen-Shannon Divergence score (B) Shannon Entropy score (C) Sum of Pairs score (D) Relative accessible surface area (E) ΔΔ Gu (F) absolute value (magnitude) of ΔΔGu. In all six sub-figures the pink violin plot represents the pathogenic scores and the beige plot represents the benign scores.

Pathogenic mutations are on average more destabilising than benign mutations (average stability: −1.1754686275, −0.250385 respectively, p = 0.0001139791). We also see a greater decrease in relative accessible surface area caused by pathogenic mutations than benign mutations (average relative accessible surface area: 0.2121568627, 0.4615 respectively, p = 1.935697E-05, **Figure 4D,E**)

We also see a negative correlation between the stability score and conservation score with all three scoring methods: Jensen Shannon divergence, Shannon Entropy and Sum of Pairs (Pearson correlation coefficient −0.378501, −0.361740 and −0.506794 respectively) and a negative correlation between relative accessible surface area and conservation score with all 3 scoring methods (Pearson correlation coefficient: −0.547688, −0.568696 and −0.488288 respectively) in this subset. Additionally, there is a positive correlation between the stability score and relative accessible surface area (Pearson correlation coefficient: 0.457250).

Taking the absolute value the average score for pathogenic mutations is 1.2617039216 and the average score for benign mutations is 0.495795 (p = 9.957418E-05). We also see a positive correlation between the stability score and conservation score with all three scoring methods: Jensen Shannon Divergence, Shannon Entropy and Sum of Pairs (Pearson correlation coefficient: 0.267980, 0.281157 and 0.458593 respectively) which suggests that the magnitude of the stability change is greater the more conserved the amino acid is. We see a negative correlation between the stability change and the relative accessible surface area, though the correlation is weaker than in the previous analysis (Pearson correlation coefficient: −0.452546). (see **Figure 4F**)

Combining all the subsets we have 50 genes containing 449 pathogenic mutations and 99 benign mutations total after eliminating amino acid deletions and mutations resulting in stop codons. We see the same trends in the conservation analysis for this subset than we see on the conservation analysis performed in the entire dataset: throughout all methods pathogenic mutations are more conserved than benign mutations (Jensen Shannon Divergence average: 0.7753923438, 0.6251661616 respectively p = 2.80E-15, Shannon Entropy average: 0.739741183, 0.4003717172 respectively p = 1.62E-23, Sum of Pairs average: 4.0318704688, 1.3970077778 p = 5.28E-24). Note: given that the current protein analysis only includes pathogenic and benign mutation scores, figures and data comparing pathogenic mutations versus the average amino acid in the protein are not included (see **Figure 5A-C**).

**Figure 5:**
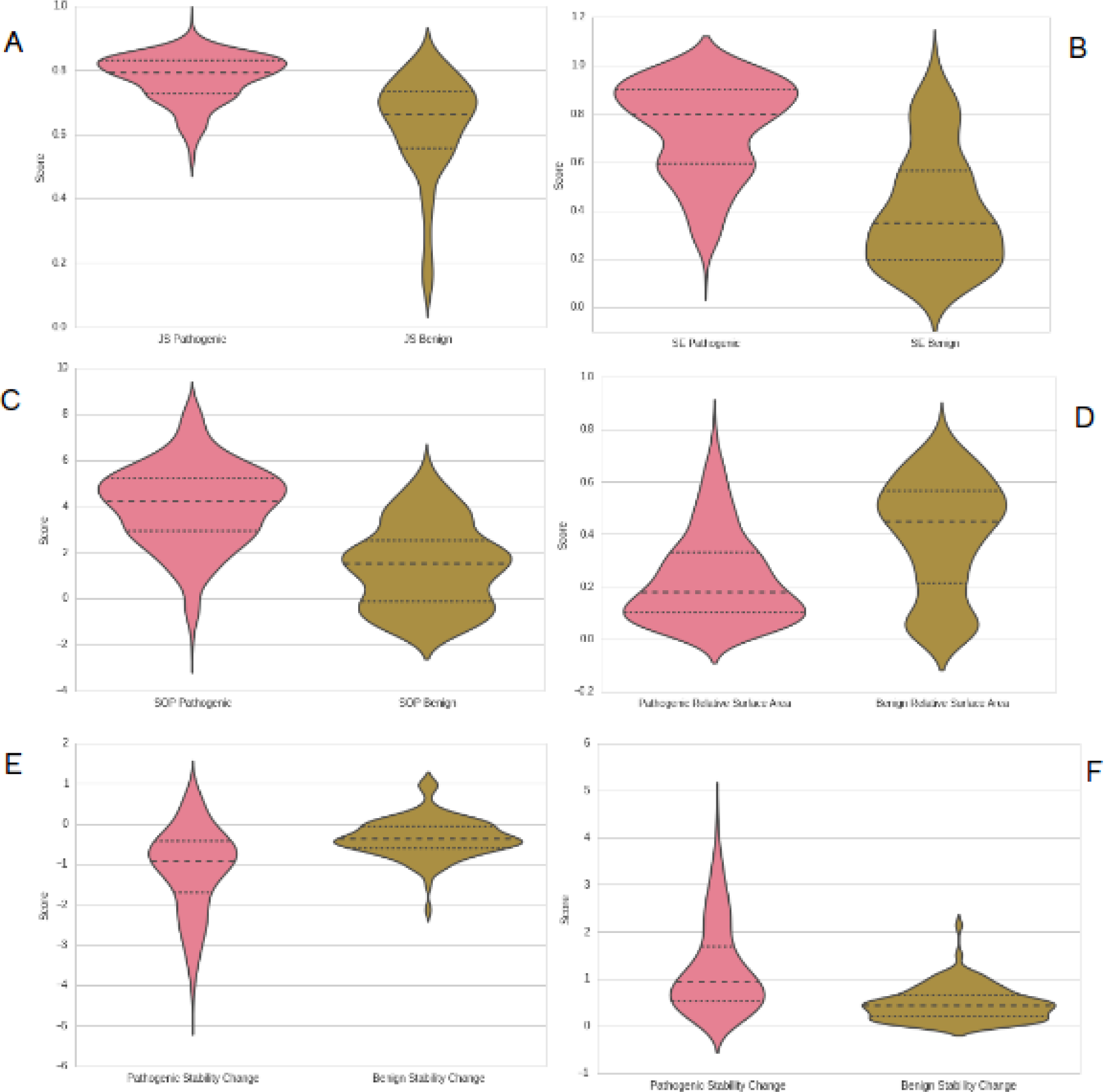
violin plots for the intersection of the 8 PANTHER subsets showing (A) Jensen-Shannon Divergence score (B) Shannon Entropy score (C) Sum of Pairs score (D) Relative accessible surface area (E) ΔΔGu (F) absolute value (magnitude) of ΔΔGu. In all six sub-figures the pink violin plot represents the pathogenic scores and the beige plot represents the benign scores.

Pathogenic mutations are on average more destabilising than benign mutations (average stability: −1.1220198218, −0.3489525253 respectively, p = 2.29E-19). We also see a greater decrease in relative accessible surface area caused by pathogenic mutations than benign mutations (average relative accessible surface area: 0.22922049, 0.3953535354 respectively, p = 8.05E-11). (See **Figure 5D,E**)

While some of the subsets have a smaller average for both stability and relative accessible surface area for pathogenic mutations, the difference between the average stability and relative accessible surface for the pathogenic mutations in the union and the individual subsets isn’t statistically significant.

We also see a negative correlation between the stability score and conservation score with all three scoring methods: Jensen Shannon divergence, Shannon Entropy and Sum of Pairs (Pearson correlation coefficient −0.358170, −0.335124 and −0.417305 respectively) and a negative correlation between relative accessible surface area and conservation score with all 3 scoring methods (Pearson correlation coefficient: −0.508771, −0.535125 and −0.492871 respectively) in this subset. Additionally, there is a positive correlation between the stability score and relative accessible surface area (Pearson correlation coefficient: 0.475668).

Taking the absolute value the average score for pathogenic mutations is 1.2139280624 and the average score for benign mutations is 0.4717909091 (p: 4.414428E-23). We also see a positive correlation between the stability score and conservation score with all three scoring methods: Jensen Shannon divergence, Shannon Entropy and Sum of Pairs (Pearson correlation coefficient: 0.267980, 0.281157 and 0.458593 respectively) which suggests that the magnitude of the stability change is greater the more conserved the amino acid. We see a negative correlation between the stability change and the relative accessible surface area, though the correlation is weaker than in the previous analysis (Pearson correlation coefficient: −0.452546). (see **Figure 5F**)

### Case Analysis: NPC1

After eliminating amino acid deletions and mutations resulting in stop codons we’re left with 38 pathogenic mutations and six benign mutations. We see the same trends in the NPC1 conservation analysis than we see in the conservation analysis performed in the entire dataset: across all methods pathogenic mutations are more conserved than benign mutations (Jensen Shannon Divergence average: 0.6946360526, 0.5859666667 respectively p = 0.01187779, Shannon Entropy average: 0.5632457895, 0.3298483333 respectively p = 0.008826891, Sum of Pairs average: 2.2894181579, −0.0415716667 p = 0.003876127). Note: given that the current protein analysis only includes pathogenic and benign mutation scores, figures and data comparing pathogenic mutations versus the average amino acid in the protein are not included (**Figure 6A-C**).

**Figure 6:**
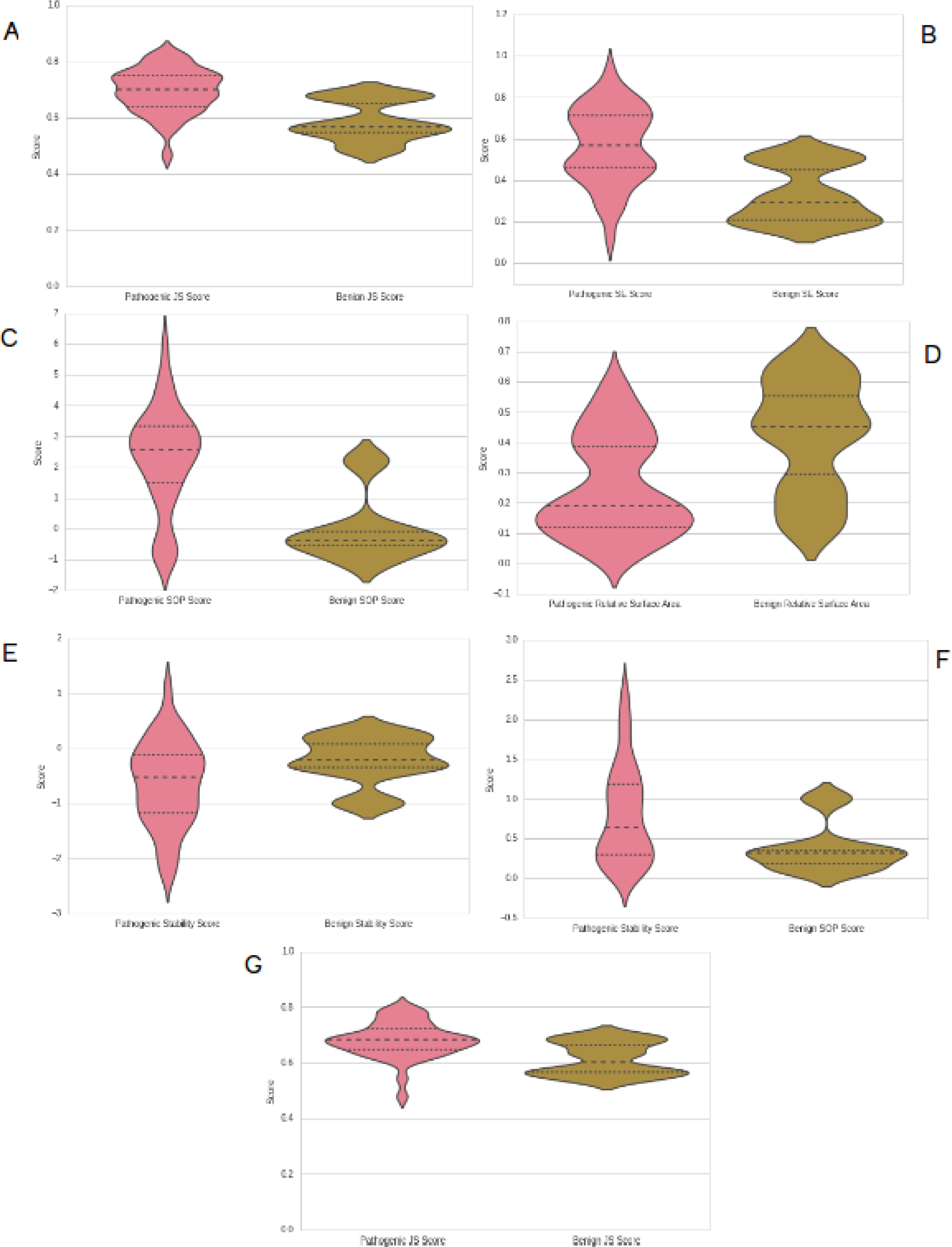
violin plots for ClinVar NPC1 data showing (A) Jensen-Shannon Divergence score (B)Shannon Entropy score (C) Sum of Pairs score (D) Relative accessible surface area (E) ΔΔGu (F) absolute value (magnitude) of ΔΔGu (G) Jensen-Shannon divergence scores including a 7 amino acid window. In all seven sub-figures the pink violin plot represents the pathogenic scores and the beige plot represents the benign scores.

**Figure 7:**
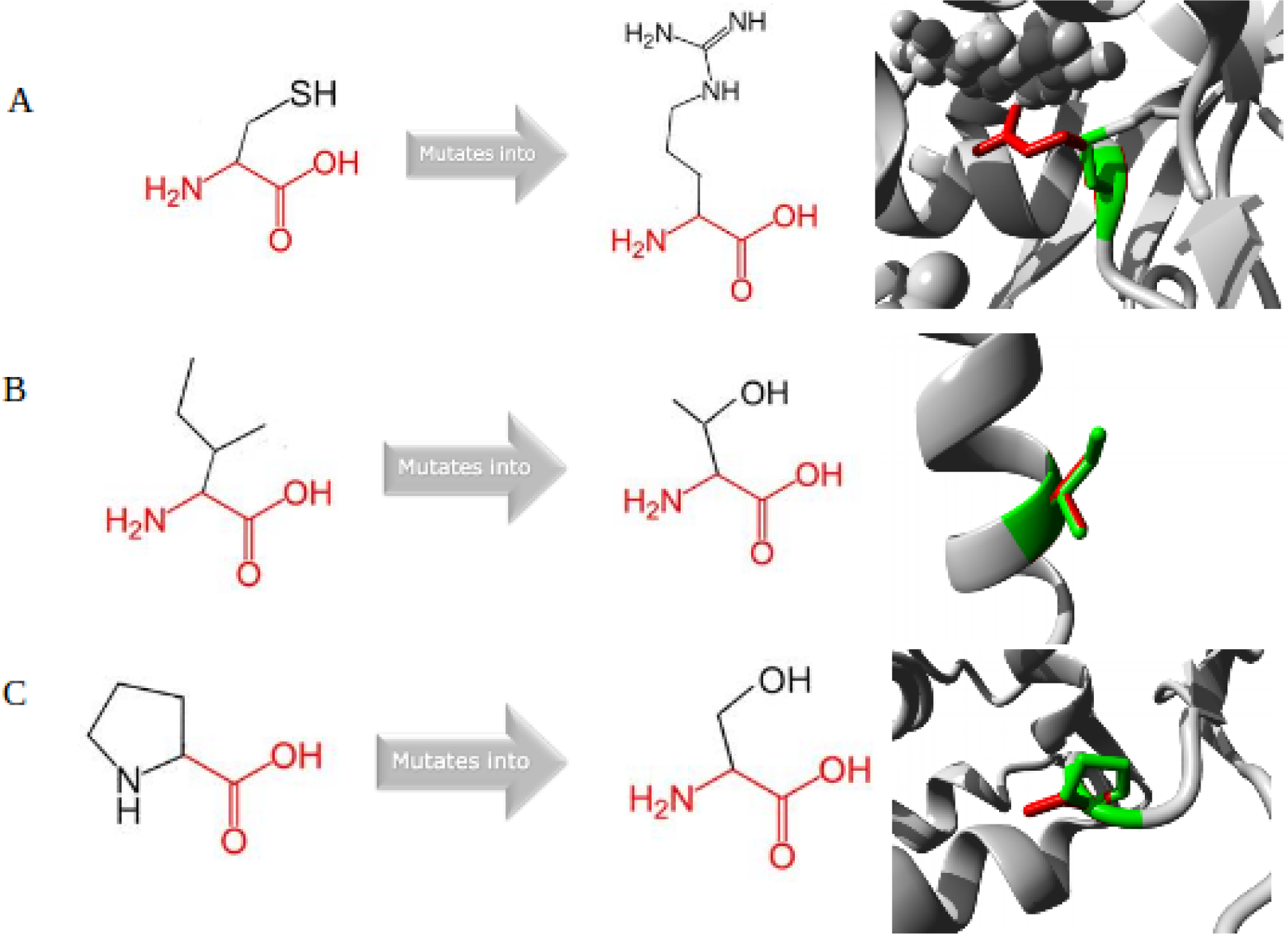
visualization of the amino acid change as well as the mutated and benign residues on the protein structure as seen on the HOPE reports. (A)C113R, (B) I1061T, (C) P434S

**Figure 8:**
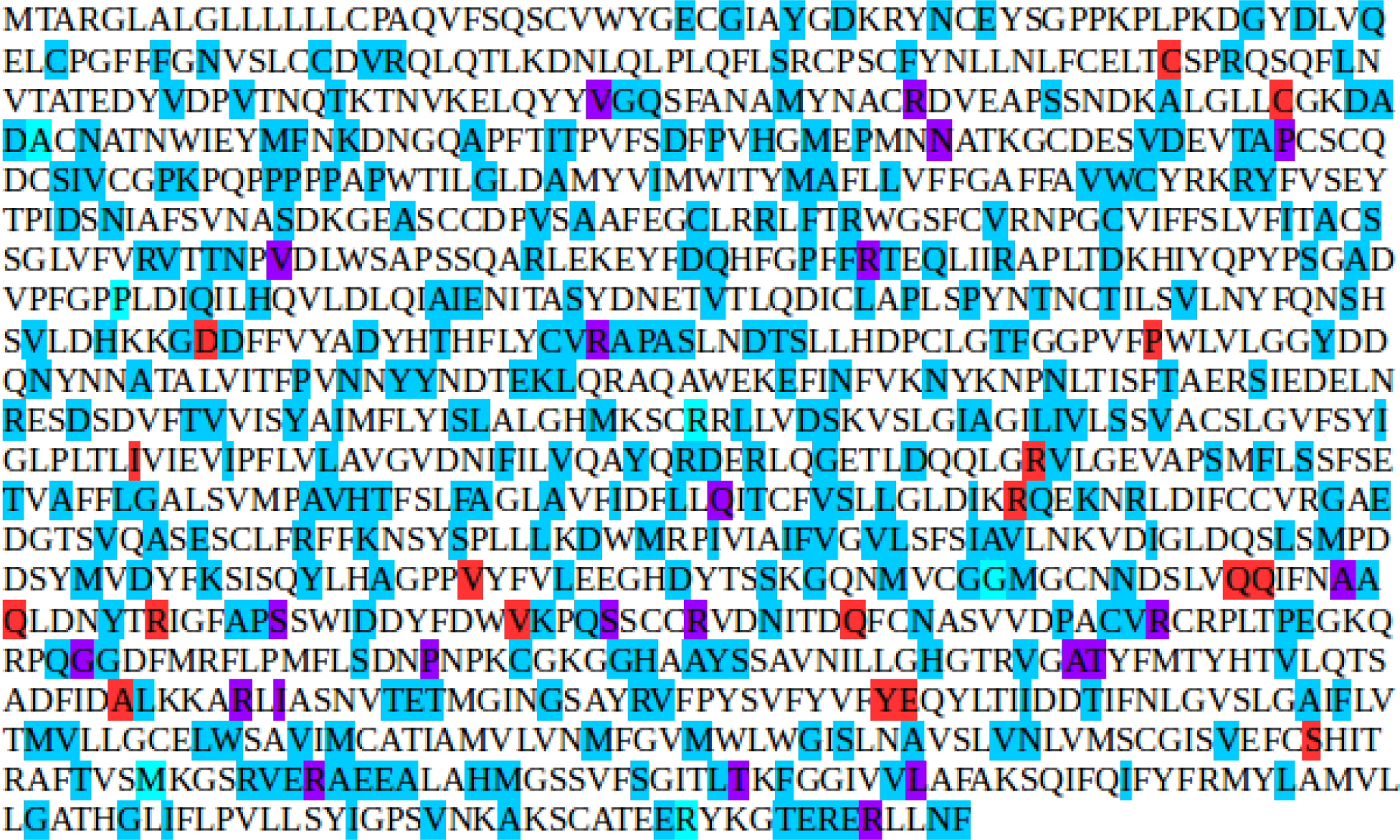
NPC1 human sequence showing pathogenic mutations (highlighted in red), ExAC mutations (highlighted in blue) and benign mutations (highlighted in cyan). Purple mutations indicatet he mutation was in both the pathogenic and ExAC set. All ClinVar benign mutations were also in the ExAC set.

Pathogenic mutations are on average more destabilising than benign mutations (average stability: −0.6427315789, −0.22485 respectively, p = 0.06223394). We also see a greater decrease in relative accessible surface area caused by pathogenic mutations than benign mutations (average relative accessible surface area: 0.2544736842, 0.42 respectively, p = 0.05569865). Note that while the difference is not statistically significant we do see the same trend as in the other subsets (**Figure 6D,E**).

We also see a negative correlation between the stability score and conservation score with all three scoring methods: Jensen Shannon divergence, Shannon Entropy and Sum of Pairs (Pearson correlation coefficient −0.382738, −0.398660 and −0.450580 respectively) and a negative correlation between relative accessible surface area and conservation score with all three scoring methods (Pearson correlation coefficient: −0.540824, −0.608378 and −0.502483 respectively) in this subset. Additionally, there is a positive correlation between the stability score and relative accessible surface area (Pearson correlation coefficient: 0.355580).

Taking the absolute value the average score for pathogenic mutations is 0.7982210526 and the average score for benign mutations is 0.37015. Unlike in the analysis above (which doesn’t take the absolute value) this difference is statistically significant (p = 0.0253726). We also see a positive correlation between the stability score and conservation score with all three scoring methods: Jensen Shannon divergence, Shannon Entropy and Sum of Pairs (Pearson correlation coefficient: 0.424044, 0.440968 and 0.504515 respectively) which suggests that the magnitude of the stability change is greater the more conserved the amino acid is. We see a negative correlation between the stability change and the relative accessible surface area, though the correlation is weaker than in the previous analysis (Pearson correlation coefficient: −0.286170, **Figure 6F**)

Given that Capra and Singh reported that incorporating the conservation of adjacent amino acid positions lead to an increase in performance of several conservation measures’ ability to predict functional sites, we decided to redo the conservation analysis using a window of size seven (three amino acids on either side of the amino acid of interest). For Jensen Shannon Divergence we see a weaker signal than we saw in the analysis before: the average score for pathogenic mutations is 0.6798907143, for benign mutations it is 0.6135683333 and the p value is 0.03131151 (**Figure 6G**)

To further assess what these scores could tell us about the expected phenotype produced by a mutation we summed the three conservation scores and the stability score (after taking the absolute value) and ranked the mutations from highest to lowest score in order to examine their characteristics in more depth. The ten highest-scoring mutations (in order) are C113R, Y1088C, V889M, P1007A, R1186H, R789H, T1205K, V378A, R404Q and I1061T. The highest scoring benign mutation is P434S, which is the position 20th of the ranked list. The average score for pathogenic mutations is 4.3455, and 1.244 for benign mutations (standard deviation: 2.345 and 1.611 respectively, p = 0.00437833). All of the aforementioned mutations were analyzed using HOPE^41^: an online tool that collects structural data from several sources, including 3D protein structures, UniProt annotations, and Reprof predictions and then produces a report detailing the predicted effect of any given protein mutation. Three sample outputs are shown below, the rest can be found on **Supplementary Materials/Figures**.

### C113R

The mutant residue is bigger than the wild-type residue. The wild-type residue was neutrally charged while the mutant one is positively charged. Additionally, the wild-type residue is more hydrophobic than the mutant one and hence it might cause loss of hydrophobic interactions in the core of the protein. Based on the 3D structure the residue is involved in a cysteine bridge, which is important for stability. While the mutated residue is not in direct contact with a ligand it could affect the local stability and therefore affect the ligand contacts made by the nearby residues. The mutation is located in a lumenal domain, associated with NPC. Given that neither the mutant residue nor a residue with similar properties was observed at this position in homologous sequences this mutation is probably damaging. It is located near a highly conserved position. BLOSUM score: -3

### I1061T

The mutant residue is smaller than the wild-type residue. The wild-type residue is more hydrophobic than the mutant which might cause a loss of hydrophobic interactions in the core of the protein. The mutation is located in a lumenal domain, associated with NPC. BLOSUM score: −1

### P434S

The mutant residue is smaller than the wild-type residue and the wild-type is more hydrophobic than the mutant residue. The mutation is located within a lumenal domain. The wild-type amino acid is a proline, since proline is known to be very rigid residue and therefore induce a special backbone conformation that the mutation might disturb. The wild-type amino acid is very conserved but a few other residue types have been observed at this position too, based on conservation scores the mutation is probably damaging. BLOSUM score: −1

### Incorporating ExAC data

To expand our mutation set we then looked at missense mutations in ExAC^42^^,^^43^, a dataset which contains exome DNA sequence data for 60,706 unrelated individuals with diverse ancestries and is released under a Fort Lauderdale Agreement. We found 424 missense mutations, although a few of them didn’t align, bringing down the total to 411. We then ran the same conservation and protein analyses as described above.

On average the set of ExAC mutations is less conserved than the average amino acid in the protein (JS divergence average: 0.638197912621359, 0.648706048513302, respectively, p: 0.0187427853018044; Shannon Entropy average: 0.466399467918623, 0.445493155339806, p: 0.0181423791777184; sum of pairs average: 1.66869084507042, 1.47736861650485, p: 0.014838270283577), consistent with the hypothesis that variants in a random sample are more likely to be benign than pathogenic and hence occur in less conserved residues. However, even though the average is lower we see a greater range of scores than in benign mutations (see **Figure 9A-C**)

**Figure 9:**
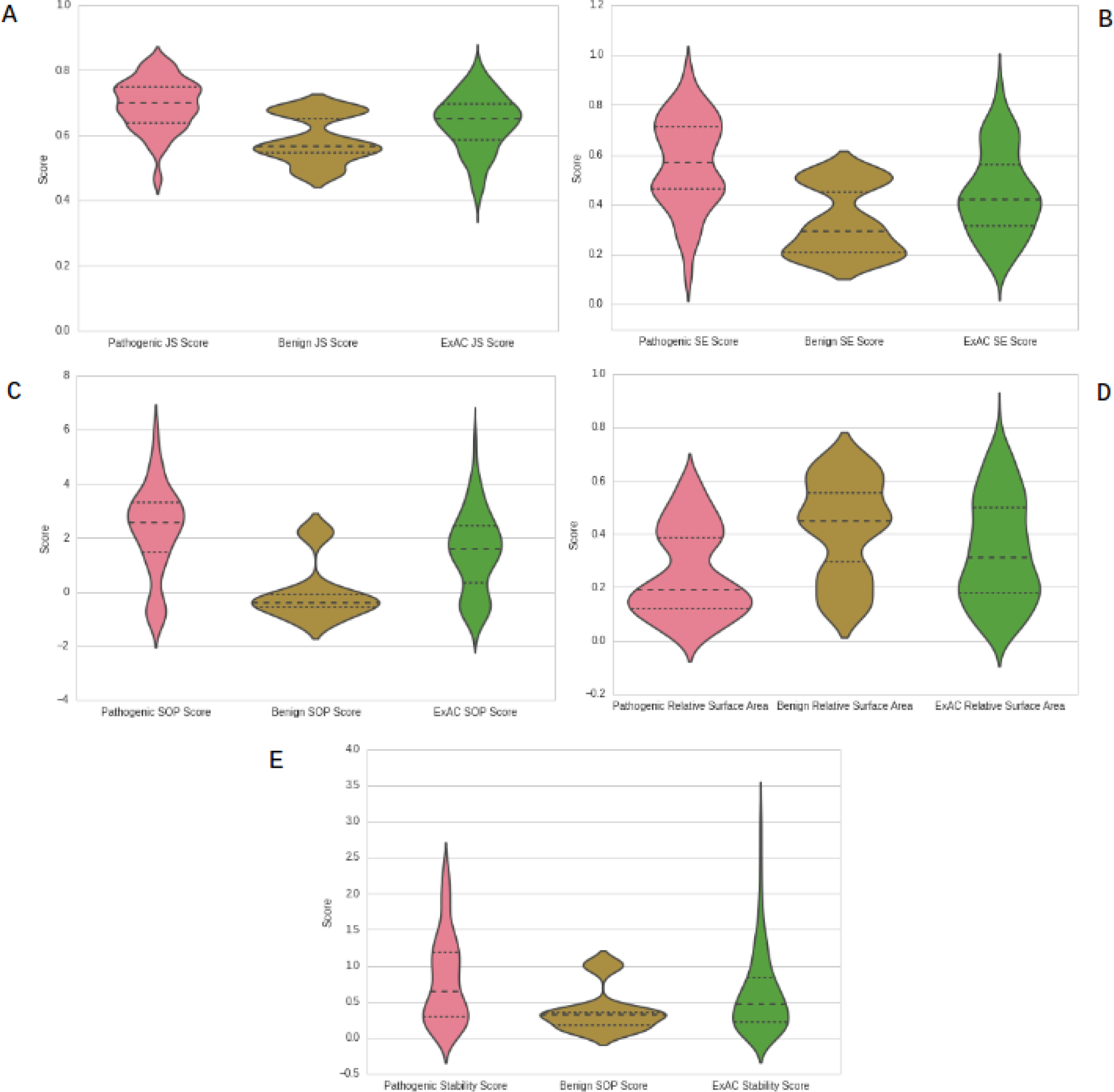
violin plots for NPC1 data showing (A) Jensen-Shannon Divergence score (B) Shannon Entropy score (C) Sum of Pairs score (D) Relative accessible surface area (E) absolute value(magnitude) of ΔΔGu. In all five sub-figures the pink violin plot represents the pathogenic scores, the beige plot represents the benign scores and the green plot represents the ExAC scores.

Taking the absolute value of the stability score the average score for ExAC mutations is 0.623086618 compared to 0.7982210526 for pathogenic mutations and 0.37015 for benign mutations, p: 0.05080188 and 2.918929E-08 respectively. On average there is a greater decrease in relative accessible surface area caused by ExAC mutations than pathogenic mutations but not benign mutations (average relative accessible surface area: 0.3337712895, 0.2544736842, 0.42 respectively, p value: 0.004318451 and 0.007448051 respectively). We also see a positive correlation between the stability score and conservation score with all 3 scoring methods: jensen shannon divergence, shannon entropy and sum of pairs (Pearson correlation coefficient: 0.325908, 0.320205 and 0.333345 respectively) which suggests that the magnitude of the stability change is greater the more conserved the amino acid is. We see a negative correlation between the stability change and the relative accessible surface area (Pearson correlation coefficient: −0.358528). Note that all but the last correlation are weaker than on the Clinvar analysis. (See **Figure 9D,E**)

Comparing to all of the mutations in the analysis by subsets the set of ExAC mutations is on average less conserved than pathogenic mutations but more conserved than benign mutations (JS divergence average: 0.6381979126, 0.7753923438, 0.6251661616 respectively, p: 3.899738E-86, 0.2152776 respectively; Shannon Entropy average: 0.4454931553, 0.739741183, 0.4003717172 p: 4.648408E-77, 0.04094601; sum of pairs average: 1.4773686165, 4.0318704688, 1.3970077778 p: 1.596554E-75, 0.3397373) (see **Figure 10A-C**)

**Figure 10:**
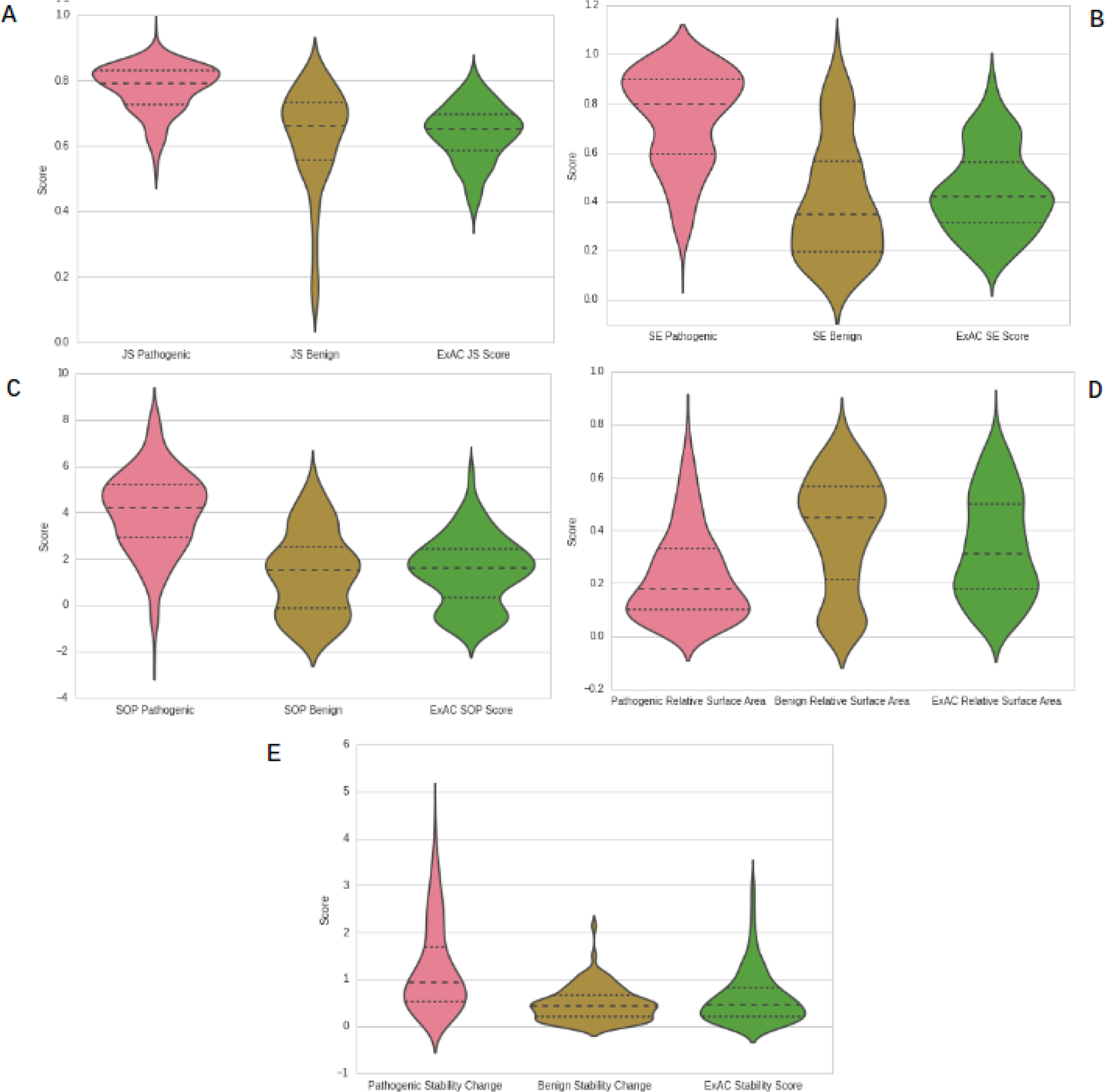
violin plots comparing subsets and ExAC data showing (A) Jensen-Shannon Divergence score (B) Shannon Entropy score (C) Sum of Pairs score (D) Relative accessible surface area (E) absolute value (magnitude) of ΔΔGu. In all five sub-figures the pink violin plot represents the pathogenic scores, the beige plot represents the benign scores and the green plot represents the ExAC scores.

**Figure 11:**
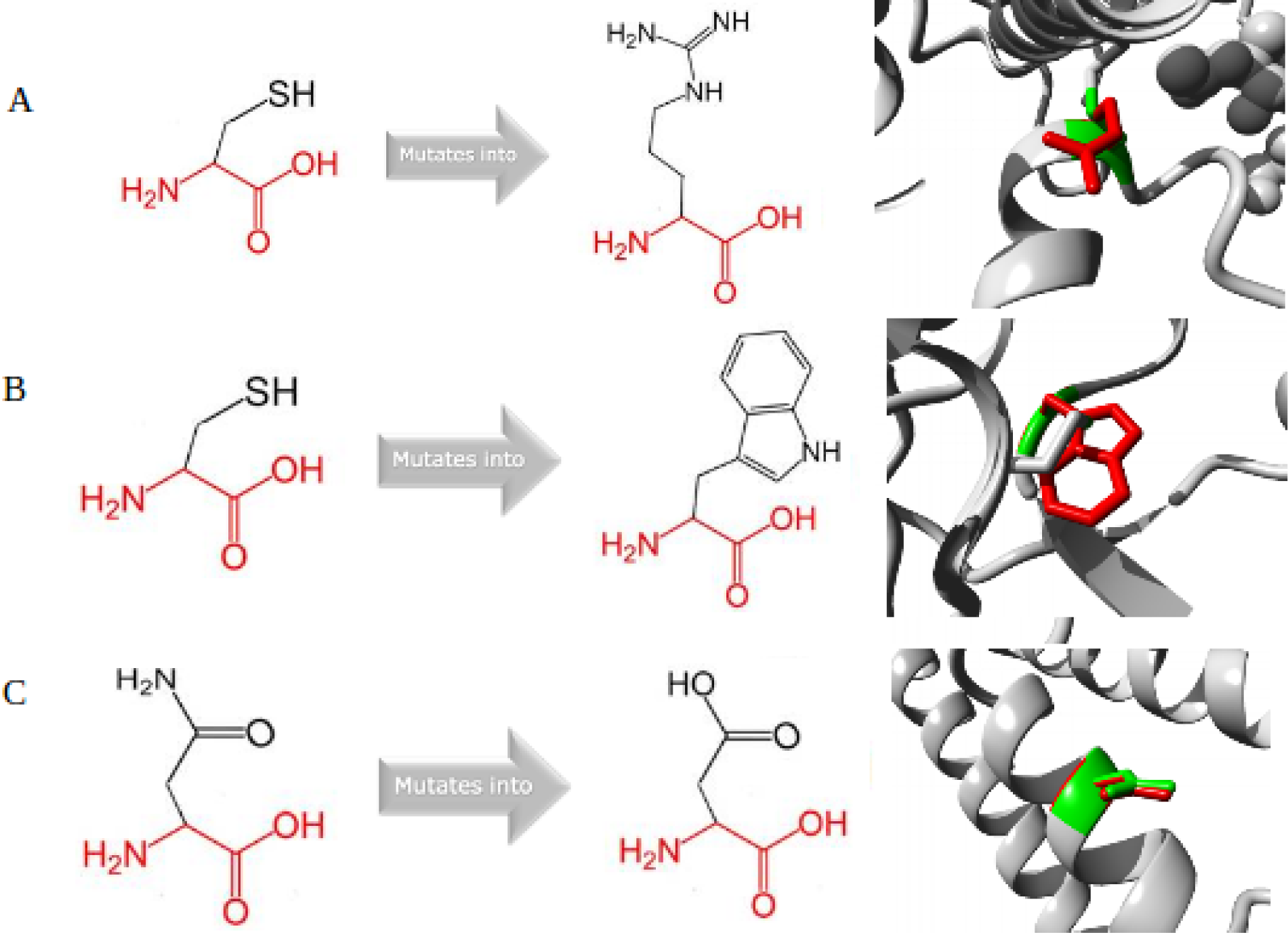
visualization of the amino acid change as well as the mutated and benign residues on the protein structure as seen on the HOPE reports. (A) C63R, (B) C75W, (C) N1156D

**Figure 12:**
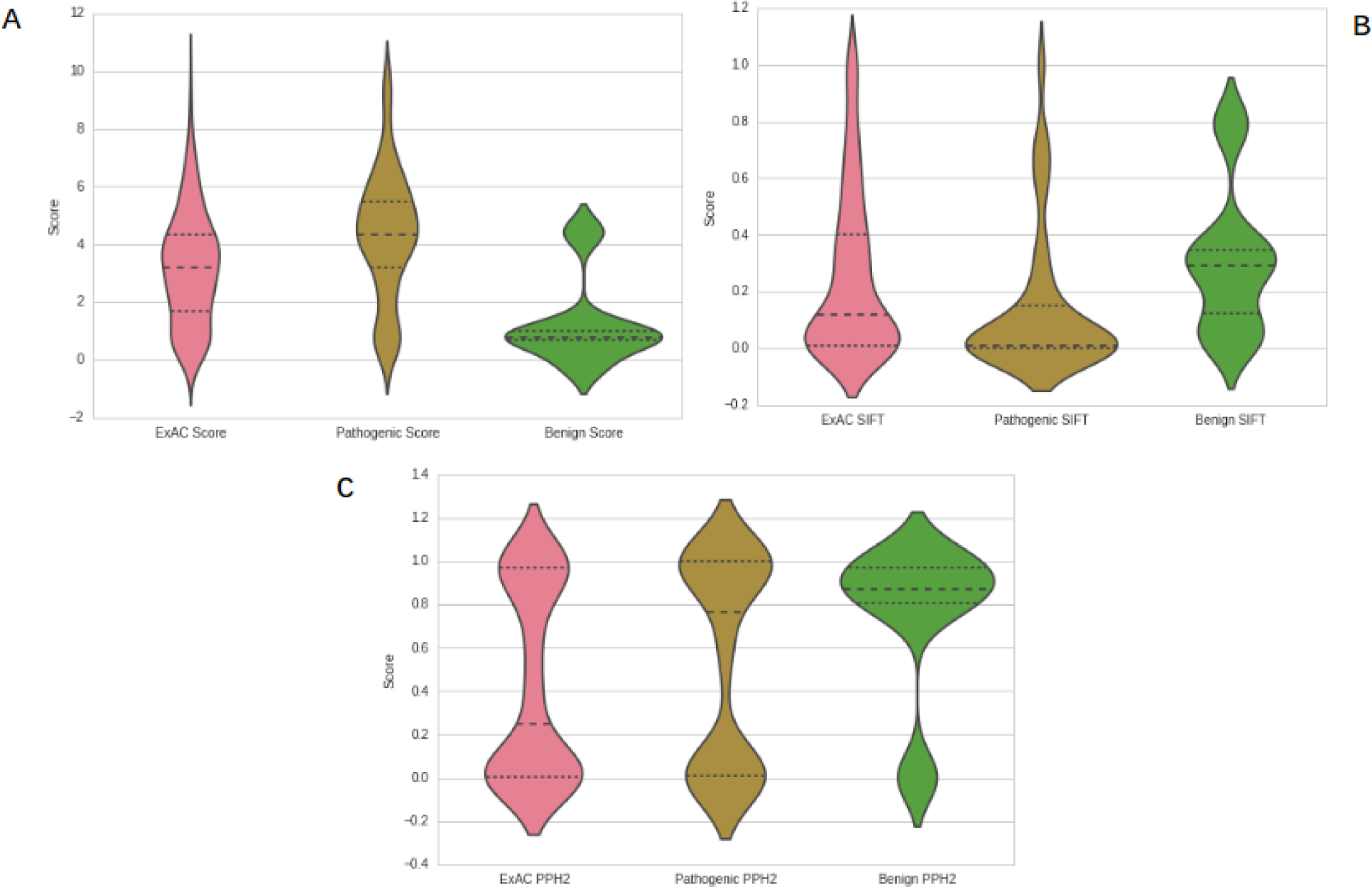
violin plots showing (A) scores using conservation and the absolute value of the stability change, (B) SIFT scores and (C) PolyPhen2 scores. In all three sub-figures the pink violin plot represents ExAC scores, the beige plot represents the Pathogenic scores and the green plot represents the benign scores.

Taking the absolute value of the stability score the average score for ExAC mutations is 0.623086618 compared to 1.2139280624 for pathogenic mutations and 0.4717909091 for benign mutations, p: 1.323672E-25 and 0.0007906132 respectively. On average there is a greater decrease in relative accessible surface area caused by ExAC mutations than pathogenic mutations but not benign mutations (average relative accessible surface area: 0.3337712895, 0.22922049, 0.3953535354 respectively, p value: 7.900523E-16 and 0.005646136 respectively). We also see a positive correlation between the stability score and conservation score with all 3 scoring methods: Jensen Shannon Divergence, Shannon Entropy and Sum of Pairs (Pearson correlation coefficient: 0.407386, 0.403053 and 0.460415 respectively) which suggests that the magnitude of the stability change is greater the more conserved the amino acid is. We see a negative correlation between the stability change and the relative accessible surface area (Pearson correlation coefficient: −0.447097). Note that all but the last correlation are stronger than on the subsets analysis. (See **Figure 10D,E**)

The mutations were scored the same way as before (with the sum of the three conservation scores and the stability score). The average score for this dataset is 3.1766304866 and the standard deviation is 1.9588718594. Two mutations have a score higher than three standard deviations above the mean (C976G and C63R) and fifteen have a score higher than two standard deviations above the mean (C976G, C63R, C1011S, C75W, F763S, F842S, N1156S, N1156D, D944N, Y1019C, F537S, V1165M, F703I, R1186C, C516F). We proceeded to analyse these mutations using HOPE to gain a better understanding of their possible functional effect. Three sample outputs are shown below, the rest can be found on **Supplementary Materials/Figures**.

### C63R

The mutant residue is bigger than the wild-type. The wild-type is neutral, the mutant is positively charged which can lead to protein folding problems. The wild-type is more hydrophobic than the mutant. The wild-type is involved in a cysteine bridge which is important for the stability of the protein, therefore the mutation can cause a loss of this interaction and destabilization of the structure. The mutation is located within a lumenal domain. The variant is pathogenic. Mutagenesis experiments have been performed on this position and mutation of the wild-type residue into S has a loss of function effect. The residue is 100% conserved. BLOSUM score: -3

### C75W

The mutant residue is bigger than the wild-type. The mutation is located within a lumenal domain. Only this residue type was found at this position. Mutation of a 100% conserved residue is usually damaging.

### N1156D

The wild-type residue was neutral, the mutant is negatively charged. The residue is located in a transmembrane domain. Mutations to both isoleucine and serine were found in this position, both are pathogenic and associated with NPC. The mutated residue is located in a protein patched/dispatched domain which is important for the protein’s activity and in contact with residues in another domain, the mutation might affect this interaction and therefore protein function.

### Benchmarking

To see how our scoring compared to other functional prediction tools we scored both the Clinvar and the ExAC mutations in NPC1 with both SIFT and PolyPhen2. Out of the 15 ExAC mutations listed above, SIFT predicted all 15 of them to be damaging (SIFT score < 0.05), while Polyphen2 predicted 11 arere probably damaging and 4 (C1011S, F842S, N1156S, F703I) are benign (from above we can see that out of those 4, N1156S has been observed to be pathogenic).

In the Clinvar data we have a total of 38 pathogenic mutations and six benign mutations. Using our scoring method two pathogenic mutations are more than two standard deviations above the mean; four are more than one standard deviation above the mean; 19 pathogenic mutations and one benign mutation are less than one standard deviation above the mean; seven pathogenic mutations are less than one standard deviation below the mean; and six pathogenic and five benign mutations are more than one standard deviation below the mean. SIFT predicted 25 pathogenic mutations to be damaging, 13 pathogenic mutations to be tolerated, one benign mutation to be damaging and give benign mutations to be tolerated. Polyphen2 predicted 16 pathogenic mutations to be probably damaging, six pathogenic mutations to be possibly damaging, 16 pathogenic mutations to be benign, two benign mutations to be probably damaging, three benign mutations to be possibly damaging and one benign mutation to be benign. This suggests our scoring method is comparable to SIFT and potentially more accurate than Polyphen2, at least in the case of NPC1.

The images above are plots of the following: the score we computed as explained above, SIFT scores, where scores less than or equal to 0.05 are damaging and above 0.05 are tolerated, and Polyphen2 scores, where the score indicates the probability of a mutation being damaging. We also see weak correlations between the different scoring methods: our scoring method and SIFT are negatively correlated, as expected, with a Pearson Correlation Coefficient of −0.309901. Our scoring method and Polyphen2 are positively correlated, with a Pearson Correlation Coefficient of 0.269999. SIFT and Polyphen2 are negatively correlated, with a Pearson Correlation Coefficient of −0.365460.

## Discussion

Throughout this paper we have reported a number of findings. First, in a pilot set of 260 Mendelian disease genes affecting cellular organelles, amino acids targetted by pathogenic mutations are significantly more conserved than the average amino acid in the protein and significantly more conserved than amino acids altered by benign mutations. This result not only supports conservation being a powerful component for predicting residues that might be important for Mendelian disease research, but also provides further support for the use of non-mammalian and even invertebrate animals as Mendelian disease models. Pathogenic mutations occurring at more conserved amino acid residues suggests that these amino acids are functionally important. We are aware this is not the first time mutation conservation has been analyzed, but as is evident from Capra and Singh^3^, the context in which conservation is used can affect whether or not conservation is sufficient for predicting functional importance. To our knowledge, this study is the first time mutation conservation has been analyzed in the context of Mendelian diseases affecting cellular organelles, and given that our dataset has a small sample size compared to the total number of genes associated with Mendelian diseases, the results might not hold true with a broader and less specific sample size.

Second, after separating the 260 Mendelian disease genes into subsets according to their function, we observed the same trends we saw in the entire dataset. The subsets themselves contain a small number of genes, and we only analyzed a small set of the subsets generated by PANTHER. However, the fact that the results reproduce across different subsets suggest that the results might hold in a broader set of Mendelian diseases. Additionally, statistical significance was not reached in a few of the subsets and individual genes, but the negative correlation between the number of mutations and the p value (as seen in **Figure 2**) suggest that a stronger, rather than weaker, signal would be seen if more genes and mutations were added to the dataset.

Third, our results combined with the results of Folkman *et al^35^* and Casadio *et al^36^* suggest that the magnitude of computationally predicted protein stability change upon mutation is a powerful tool for predicting functional importance that can be used alongside conservation scores for better prediction power. Neither SIFT nor PolyPhen2 uses computed ΔΔGu as part of their prediction methods, but given that EASE-MM is an entirely sequence based tool, this could be a powerful supplement to existing methods, especially for cases where the protein structure is not available.

Our analysis has several shortcomings, which we hope we can address in the near future. The analysis we presented here can easily be extended to any other genes for which homologous sequences are known, and more species aside from the six used here can be included as well. The accuracy of our predictions and how they compare to those made using other methods (like SIFT and PolyPhen2) or the broader significance (or lack thereof) that our results might have for modelling Mendelian diseases using simple organisms cannot be assessed until our results are verified in vivo. Additionally, unlike other methods, the steps involved in our analysis aren’t fully automated and require a substantial amount of human input. We should note that while making our analysis easier to reproduce and perform by others is something we are interested on working on, the analysis presented above is not intended to be used as a standalone method to decide which mutations to use in model organisms or to predict clinical significance. Instead, it is meant to serve as an aid to help narrow down the list of candidate mutations. Mendelian diseases are very biologically complex.

Finally, all of the data included in this analysis is available on github (github.com/materechm/plabData), and all of the data used to perform the analysis came from publicly available databases. In the spirit of open science, especially as it applies to the rare disease space, we hope that this initial study sparks follow-on collaborations to expand the analysis to include a larger number of Mendelian disease genes, and to test predictions for variants of unknown significance.

